# Novel strategies to deal with salinity in *Sorghum bicolor* seedlings

**DOI:** 10.1101/2025.10.02.680130

**Authors:** Moira R. Sutka, Pablo D. Cáceres, Marina Recchi, Andrea S. Dengis, Milena E. Manzur

## Abstract

Sorghum is a crop that has become more relevant in recent years due to its uses and properties (biofuel, gluten-free flours) as well as its versatility to grow in unfavorable environmental conditions. Salinity is one of the main abiotic stresses affecting crop production and yield worldwide. The aim of this work was to study the response of sorghum seedlings to soil salinity in two genotypes with known performance to cope with water stress. Physiological parameters related to plant water status as well as the sodium content in the plant were analyzed. We studied the possible role of pricklet and microhairs in the response to salt conditions. Our results showed a differential response to salinity, probably denoting different mechanisms that involve internal water redistribution (in 200 mM NaCl) and a specific replacement of silicon by sodium (when the NaCl reaches 300 mM). The main result was that sodium was absent in all analyzed hairs and leaf surface. Surprisingly, we detected the presence of silicon inside the pricklet at 300 mM NaCl after 24 hours, but not in the microhair. NIPs aquaporins could be involucrate in silicon transport. Our novel results provide further evidence regarding the role of silicon in the response to salt stress.

**Highlight:** Grain sorghum has a different strategy to deal with salinity stress depending on the salt concentration, that involves the leaves pricklets and the migration of silicon.

## Introduction

During their life cycle, plants are exposed to a series of unfavorable environmental conditions, such as drought, flooding, and salinity that generate stress and negatively affect their growth and development. These conditions represent a challenge for multiple crops that can cope using different mechanisms (Acosta-Motos *et al.,* 2017). Soil salinity is one of the main problems in global agriculture with a total area of salt-affected soils around 1400 million ha, which represents almost the 11% of the total land area and it is in expansion at a rate of 2 million ha per year due to climate changes (Abbas *et al.,* 2013; Global status of salt-affected soils, 2024). It is very well studied that soil salinity is one of the most important causes of yield and economic loss, with effects that can reduce crop yield up to 58% (Haj-Amor *et al.,* 2022). Therefore, it is imperative to continue researching the mechanisms and responses of crops capable of tolerating the effects of salinity in order to maintain production. In this way, sorghum is one of the target species, since it is a model cereal, and its response was studied against different types of stress. Additionally, this crop has a potential use as cellulosic biofuel, its carbon metabolism (C4) is more efficient at high environmental temperatures (Hu *et al.,* 2022; Mubarak Alqahtani, 2024), and its small genome (730Mb) (Paterson *et al.,* 2009) make it a useful tool for functional genomics approaches.

Under salt stress, tolerant seedlings of different species show a relatively high ability to recover after stress. This recovery is often associated to a more adequate carbon partition between shoots and roots and to changes in absorption, transport and re-translocation of salt (De Lacerda *et al.,* 2005). In sorghum, it was observed that sodium (Na^+^) increased its concentration in leaves and roots under high external concentrations (from 50 to 200 mM), and in leaves, also proline was accumulated (Bavei *et al.,* 2011). In proteomics studies an overexpression of proteins related to stress responses was described, and the authors identify four proteins with no apparent function, which suggests that sorghum has a differential response mechanism to salinity in relation to other species responses, as maize (Kumar Swami *et al.,* 2011; Sekhwal *et al.,* 2012).

In relation to the mechanisms that allow plants to cope with salinity, there are salt bladders, salt glands and microhairs (Ramadan and Flowers, 2004; Shabala *et al.,* 2014; Yuan *et al.,* 2016). Salt bladders can have stalk cells and their own bladder, where sodium and chloride are sequestered. After a prolonged period, they detach releasing salt into the leaf surface. These salt bladders have been well characterized in two halophytes, *Mesembryanthemum crystallinum* (Agarie *et al.,* 2007) and *Chenopodium quinoa* (Adolf *et al.,* 2013). Although the molecular mechanism by which salt accumulates inside is not yet known, there is a proposed model where certain transporters are involved (Chen *et al.,* 2017; Böhm *et al.,* 2018). In contrast, salt glands have a more complex structure than bladders and can be divided into bicellular or multicellular. Bicellular glands have one collected cell and one secretory cell, while multicellular glands can have different quantities of each (Shabala et al., 2014). The mechanism of salt transport in the glands is unclear and the pathway of sodium from the mesophyll cells to the collecting cells is still unknown. However, plasmodesmata and connections with the secretory system were found (Sager and Lee, 2014). Microhairs are structures present on the adaxial surface of the leaf. They were described in the wild rice *Porteresia coarctata* where it was shown that they accumulate and secrete NaCl (Flowers *et al.,* 1990). Recently, they were isolated and molecularly characterized (Rajakani *et al.,* 2019). In the Poaceae family, one of the most numerous among angiosperms, there are around 10000 species and 700 genera (Grass Phylogeny Working Group, 2001; 2012). Species within this family show wide variation in terms of salinity tolerance (Marcum, 2008). In *Sorghum halepense*, the presence of bicellular trichomes was described by analyzing the morphology of the leaf (McWhorter *et al.,* 1995). These authors reported secretions and positive staining reaction for callose and polysaccharides but did not detect salt crystals when plants were grown in soil containing NaCl (McWhorter *et al.,* 1995). In *S. bicolor* there is no evidence so far of the presence of structure related to salt excretion or accumulation. Like other halophyte plants, *S. bicolor* may use microhairs to extrude sodium when it is in physiological concentrations detrimental to growth. The objective of this work was to investigate the physiological responses of two genotypes of *S. bicolor* seedlings and the potential role of microhairs in sodium exclusion and/or isolation processes under salinity conditions.

## Material and methods

### Plant material and culture conditions

Experiments were performed using two sorghum [*Sorghum bicolor* (L.) Moench] inbred lines, RedLand B2 and IS9530, which are parental genotypes that originated the regularly sown lines in Argentina (Rodriguez V, pers. comm.). Plants were grown under controlled environmental conditions: 16/8 h light/dark photoperiod, light intensity of 160 ± 3 µmol m^−2^ s^−1^, 75% relative humidity and a constant temperature of 23 °C in a growth chamber. Seeds were sown in plastic containers (200 mL) filled with sterilized sand and moistened with Hoagland solution: 1.25 mM KNO_3_, 0.75 mM MgSO_4_, 1.5 mMCa(NO_3_)_2_, 0.5 mM KH_2_PO_4_, 50 μMFeEDTA, 50 μM H_3_BO_3_, 12 μM MnSO_4_, 0.70 μM CuSO_4_, 1 μM ZnSO_4_, 0.24 μM Na_2_MoO_4_, and 100 μM Na_2_SiO_3_ (Javot *et al.,* 2003). Ten days after germination seedlings were transplanted into plastic pots filled with 4.5 L of Hoagland solution and a fish tank aerator to oxygenate. To apply salt treatments, NaCl was added to the pots at 0, 100, 200 or 300 mM final concentration. Measurements were taken after 1, 4 or 24 hours of salt treatment, except for water potential that was also measured at 15 and 30 minutes. Data was collected from two to eight plants per condition, from three to eight independent experiments.

### Relative water content

Relative water content (RWC) was determined according to Turner (1981). Briefly, leaves of treated or control seedlings were cut and immediately weighed (fresh weight, FW), followed by immersing them for 24 h in distilled water to determine turgid weight (TW) and finally dried at 60 °C for 48 h to obtain dry weight (DW). Relative water content was calculated as: RWC = [(FW – DW)/(TW – DW)] × 100.

### Stomatal conductance

Stomatal conductance was measured at midday using a portable steady state diffusion leaf porometer (model SC-1, Decagon devices, Pullman, WA, USA). Measurements were taken on the abaxial surface at the center of the fully expanded leaf.

### Water potential (**Ψ**_w_)

Whole aerial part was introduced in a Scholander pressure chamber (Biocontrol Model 4, Argentina) to determine the water potential (Ψ_w_) (Scholander *et al.,* 1965). Briefly, the aerial part was cut, introduce in the Scholander chamber and close. Pressure was applied until xylem sap appeared. This pressure value is considered as water potential value that the seedling had before the cut.

### Osmotic potential (**Ψ**_osm_)

Osmotic potential in roots, leaf sheaths and leaf blades was measured as previously described by (Mahdieh *et al.,* 2008). Briefly, excised tissues from treated or control plants were placed in small columns with holes at the bottom and immediately frozen in liquid nitrogen. After thawing, the column was inserted in 1.5 mL centrifuge tube and centrifuged at 4000 x *g* for 4 min at 4 °C. Sap osmolarity of each sample was measured in a vapor pressure osmometer (Vapro 5520, Wescor, USA) and used to calculate osmotic pressure according to Van’t Hoff’s equation.

### Sodium Content

Sodium concentration was determined in roots, leaf sheaths and leaf blades of control and treated seedlings. Samples were dried at 80 °C and then subjected to acid extraction following Hunt (1982). Briefly, 0.1 g of dry tissue was transferred to a wide mouth bottle (100 mL capacity) and 50 mL of 0.5 M HCl was added. The samples were incubated in a water bath at 30 °C for 15 min. After cooling, content was filtered through Whatman paper. Sodium content determinations were made using flame emission photometry (Crudo Caamaño, Laboratorios Norte, Argentina). Calibration curve was made using serial dilutions of a standard sodium solution (326 mg Na^+^/100 mL) to calculate samples final concentrations.

### Root hydraulic conductivity

Root hydraulic conductivity measurements were carried out essentially as described by Javot *et al.,* (2003). Briefly, the whole root system of freshly detopped sorghum seedlings was immersed in a 50 mL container filled with hydroponic culture medium and inserted into a pressure chamber (Biocontrol Model 4, Argentina). The root-shoot junction was carefully threaded through the metal lid of the chamber, sealed using low-viscosity dental paste (A+ Silicone, Densell) and connected to a graduated glass micropipette. Pressure-induced sap flow (Jv) from the root system was measured at three different pressures: 0.2, 0.3 and 0.4 MPa, and was determined by collecting the exuded sap into the micropipette. After completing the measurements, the root system was removed and oven-dried to determine its dry weight. The hydraulic conductivity of an individual root system (Lpr) was calculated as the slope of the flow rate (Jv) versus pressure (P), normalized by the root dry weight.

### Microscopy images

Images were obtained from fresh seedlings using a scanning electron microscope (ESEM, FEI Quanta 250 with tungsten filament) equipped with an Energy Dispersive Spectroscopy (EDS) Ultra Dry (Thermo Fisher Scientific) that allow us to determine ions composition of samples.

### Sequence alignment

A sequence analysis was performed between aquaporins known to transport silicon in other crop species and those in sorghum, increased in saline stress to determine the degree of homology between them. Sequence search was performed using BLASTP tool (https://blast.ncbi.nlm.nih.gov/Blast.cgi) for the following NIPs aquaporin: *Glycine max* NIP2;1 (GI: 1528010474), NIP2;2 (GI: 358248754), *Oryza sativa* NIP2;1 (GI: 2447447740), NIP2;2 (GI: 2211362252), *Hordeum vulgare* NIP2;1 (GI: 296837167), NIP2;2 (GI: 383276514), *Zea mays* NIP2;1 (GI: 146325012), NIP2;2 (GI: 75308078), *Sorghum bicolor* NIP2;1 (protein ID: XP_002454286) and NIP2;2 (Protein ID XP_002438105) (Handa *et al.,* 2022; Reddy *et al.,* 2015). Protein sequences were aligned using the Clustal Omega program (Madeira *et al.,* 2024).

### Statistical analysis

Two-way analysis of variance (ANOVA) was used to analyze the effects of genotype and salt treatment on physiological seedling responses. Post-hoc Tukey’s tests were employed for mean comparisons (P ≤ 0.05). Variable normality and homogeneity of variances were previously verified in order to satisfy ANOVA’s assumptions. All statistical analyses were performed with the GraphPad Prism 8.0 software. Results are presented as mean ± SEM (standard error of the mean). Each sample size (n) is indicated in the figure legends.

## Results and discussion

### The phenotypes of both genotypes are not modified in presence of salt

The phenotype of both genotypes remained unchanged throughout the salt concentration (100 to 300 mM) and duration treatments (1, 4 and 24 h) (Fig. 1; Fig. S1). However, some physiological parameters were altered by increased salt concentration, as described in the next sections. By convention, soil is considered as saline when its electrical conductivity is at least 4 dS m^−1^, which is equivalent to 40 mM NaCl (USDA-ARS, 2008). In sorghum, plant growth begins to slow down after 6.8 dS m^−1^ (Francois *et al.,* 1984) which is equivalent to 68 mM NaCl. Unlike what has been reported for sugar beet (Vitali *et al.,* 2015), sorghum plants can maintain their turgidity throughout increases in NaCl in the growth medium (Fig. S1). Our results reinforce what is currently known about the different physiological mechanisms used by monocots and dicots to cope with salt stress, as we discuss below.

**Fig. 1.**
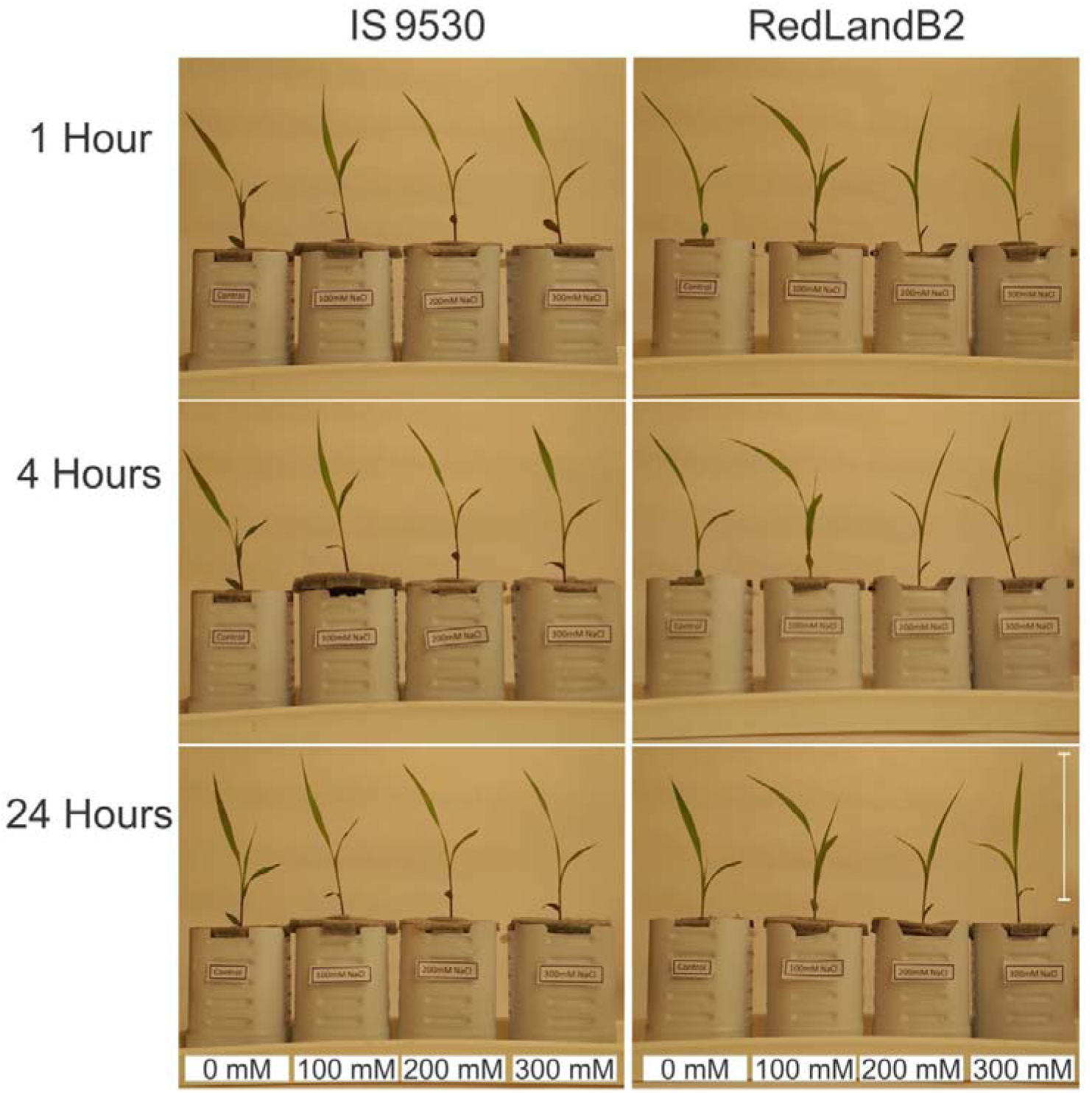
Photograph of sorghum seedlings (10 days old) of two genotypes (IS9530 and RedLand B2). Seedlings were subjected to increasing sodium concentrations: 100, 200 and 300 mM during 1, 4 and 24 hours. Scale bars= 10cm.

### Physiological parameters related to water balance are affected under salt conditions

Stomatal conductance was reduced throughout all salt treatments and all times evaluated (1, 4 and 24 hs) except at 100 mM NaCl after 24 hours (Fig. 2).

**Fig. 2.**
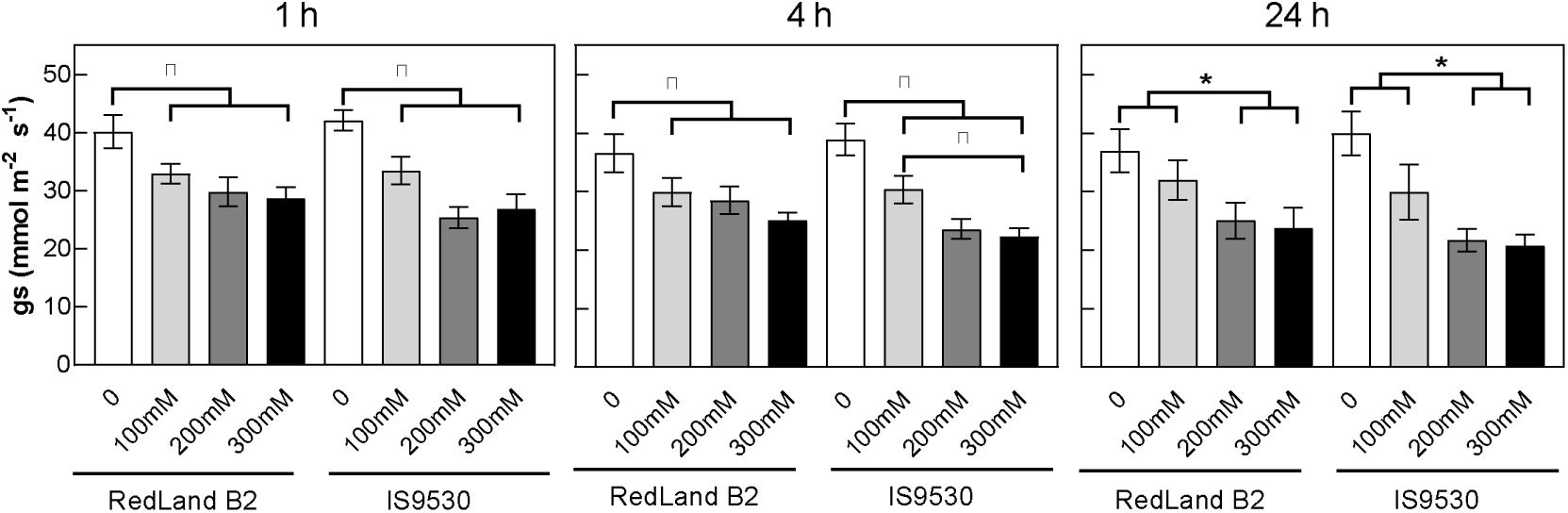
Stomatal conductance (gs, mmol m^−2^ s^−1^) measured in sorghum seedlings (10 days old) of two genotypes (RedLand B2 and IS9530). Seedlings were subjected to increasing sodium concentrations: 100, 200 and 300 mM during 1, 4 and 24 hours. Values are means ± SEM of three plants per genotype from seven independent experiment. Asterisks indicate statistically significant differences between sodium concentrations for each genotype (two-way ANOVA, Tukey post-hoc test; *P<0.05, **P<0.01, ***P<0.001, ****P<0.0001).

The hydric state of the plant is well defined by the water potential of its tissues (Turner *et al.,* 1981). Our results demonstrated that the water potential was reduced throughout all salt treatments at 1 and 4 hours. Since a linear drop in the water potential at 300 mM NaCl was not observed, one hour after the treatment, as it occurred at 100 and 200 mM, the change in water potential at 15 and 30 minutes was analyzed (Fig. 3). Surprisingly, water potential at 300 mM starts with values, as expected, lower than control (−0.3 MPa and −0.35 MPa for RedLand B2 and IS9530 respectively). However, at 24 hours only plants growing in 300 mM NaCl equalized control values in both genotypes (Fig. 3). Additionally, sheath anatomy was analyzed in order to discard a possible rupture of xilematic tissues (Fig. S2).

**Fig. 3.**
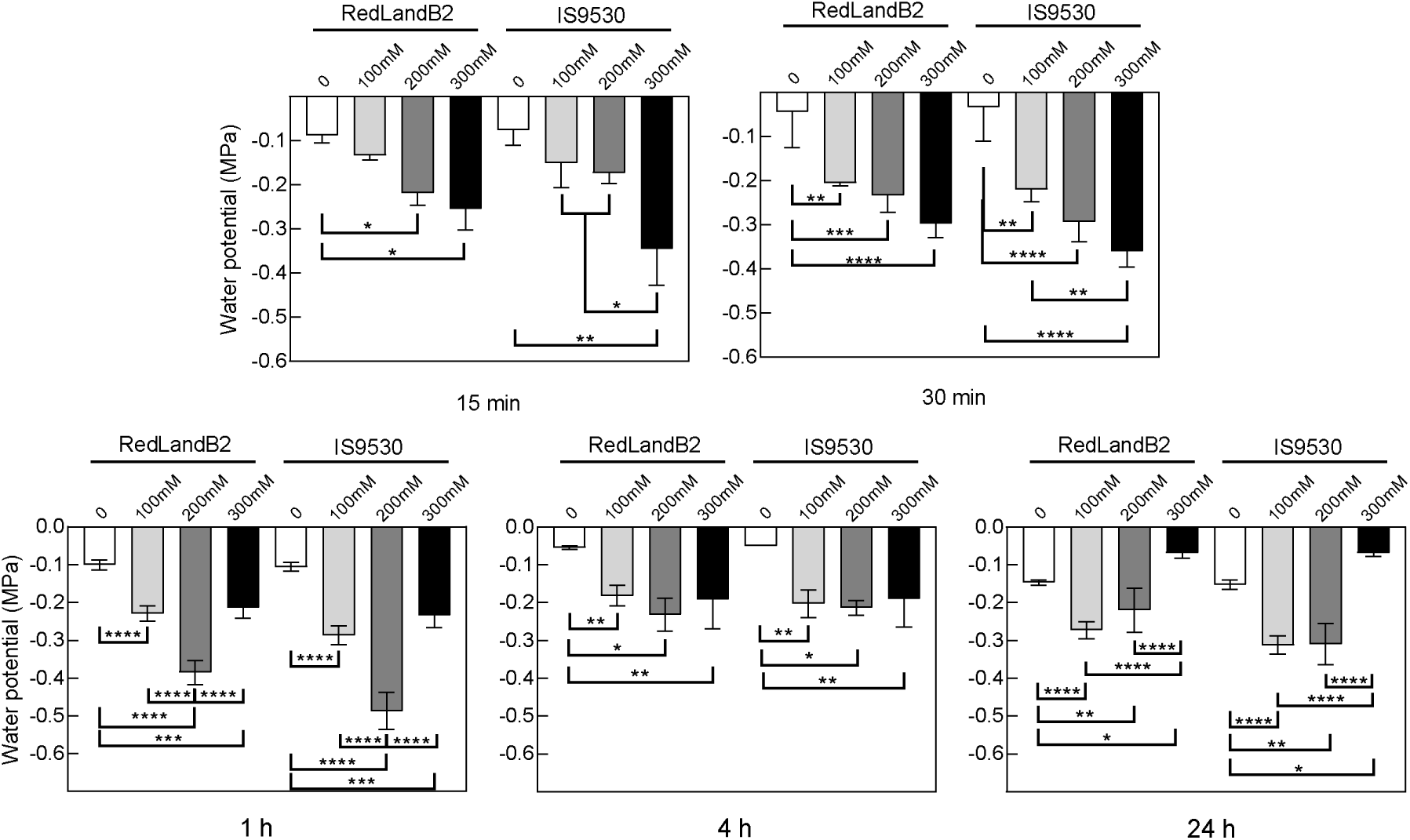
Water potential (MPa) measured in sorghum seedlings (10 days old) of two genotypes (RedLand B2 and IS9530). Seedlings were subjected to increasing sodium concentrations: 100, 200 and 300 mM during 15 and 30 minutes and 1, 4 and 24 hours. Values are means ± SEM of three plants per genotype from five or eight independent experiments. Asterisks indicate statistically significant differences between sodium concentrations for each genotype (two-way ANOVA, Tukey post-hoc test; *P<0.05, **P<0.01, ***P<0.001, ****P<0.0001).

At root level, hydraulic conductivity is reduced only under 200 mM NaCl throughout all times evaluated. At 100 and 300 mM NaCl this parameter did not change with respect to control conditions (Fig. 4).

**Fig. 4.**
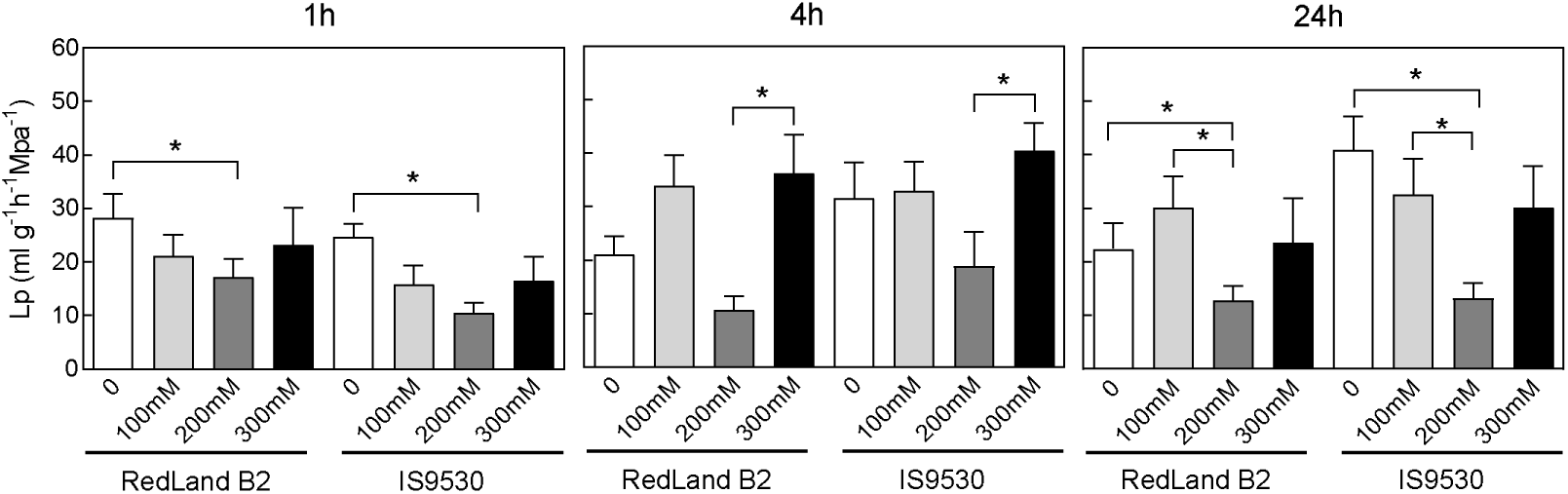
Hydraulic conductivity (ml g^−1^ h^−1^ MPa^−1^) measured in sorghum seedlings (10 days old) of two genotypes (RedLand B2 and IS9530). Seedlings were subjected to increasing sodium concentrations: 100, 200 and 300 mM during 1, 4 and 24 hours. Values are means ± SEM of nine to fourteen plants per genotype. Asterisks indicate statistically significant differences between sodium concentrations for each genotype (two-way ANOVA, Tukey post-hoc test; *P<0.05).

Finally, it is possible to affirm that the relationships between the above-mentioned parameters determine the water content of the plant. To verify this, the relative water content of both genotypes was determined for different combinations of time and NaCl concentration (Fig. 5). Our results demonstrated that relative water content was not affected at 100 mM along the time treatment, but decreased under 200 and 300 mM NaCl, after one and four hours of treatment. However, after 24 hours the RWC does not show differences with the control conditions (Fig. 5).

**Fig. 5.**
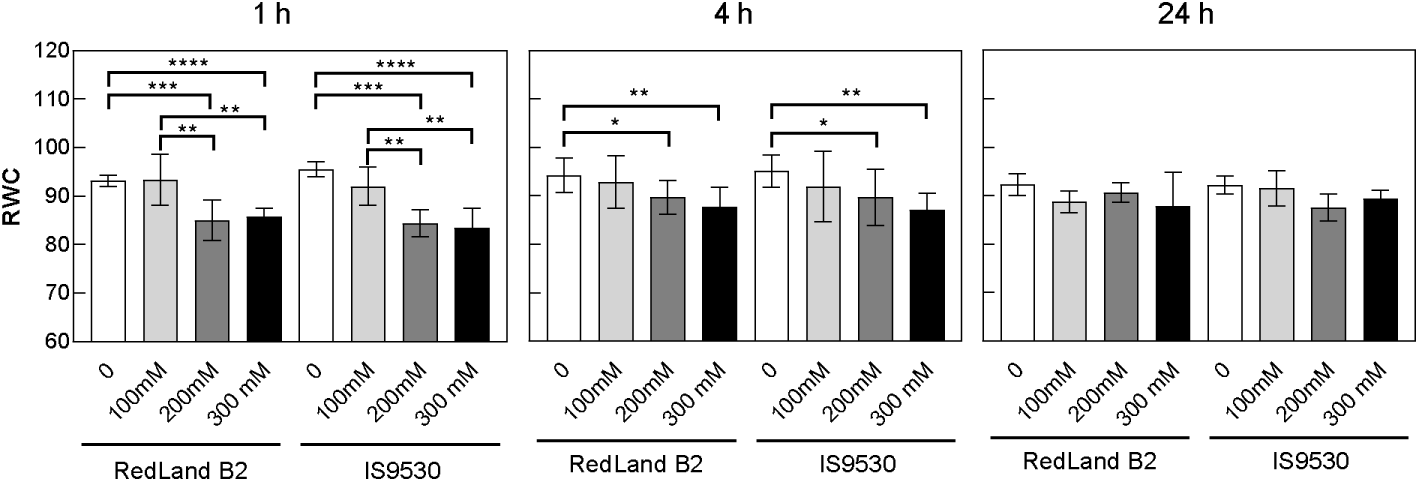
Relative water content (RWC) measured in sorghum seedlings (10 days old) of two genotypes (RedLand B2 and IS9530). Seedlings were subjected to increasing sodium concentrations: 100, 200 and 300 mM during 1, 4 and 24 hours. Values are means ± SEM of three plants per genotype from four to eight independent experiments. Asterisks indicate statistically significant differences between sodium concentrations for each genotype (two-way ANOVA, Tukey post-hoc test; *P<0.05, **P<0.01, ***P<0.001, ****P<0.0001).

In dicot salt tolerant plants, a concentration of 100 mM NaCl it is well tolerated, although *A. thaliana* is very sensitive at such concentration (Munns and Tester, 2008). The RWC recovery at 24 h, even with the stomata closed (as shown in Fig. 2), occurs in different ways for each salt concentration. At 200 mM NaCl the root hydraulic conductivity is low, indicating a water internal redistribution process, as was observed in other salt tolerant species (Lu and Fricke, 2023). This would be the first report of this type of mechanism under saline conditions in an annual crop such as sorghum. At 300 mM NaCl the mechanism could be related to the high root hydraulic capacity, allowing the water incoming trough the roots. In *A. thaliana* accession (Mr-0), showed a high root hydraulic conductivity at 100 mM NaCl (Sutka *et al.,* 2011), guarantee the optimal functionality of roots. In sorghum seedling something similar could happen, since water potential was higher than control condition, suggesting that sodium is not inside the cell. In this scenario, we analyzed the osmotic potential and sodium concentration along the entire plant.

### Sodium final location in the plant body would depend on its external concentration

At one hour of treatment, the root osmotic potential decreased in a dose-dependent manner and this pattern was conserved over time (Fig. 6). However, the osmotic potential at 300 mM cannot be explained solely by the Na^+^ concentration present in the root. However, at one hour of treatment, there was a decoupling with the response of the osmotic potential of the aerial part since it did not change significantly under any of the concentrations (Fig. 6). At 4 and 24 hours, a drop in the osmotic potential of the shoot at 200 mM and 300 mM was detected, which did correspond to the increase in Na^+^ concentration in both sheath and blade (Table 1; Fig. 6).

**Fig. 6.**
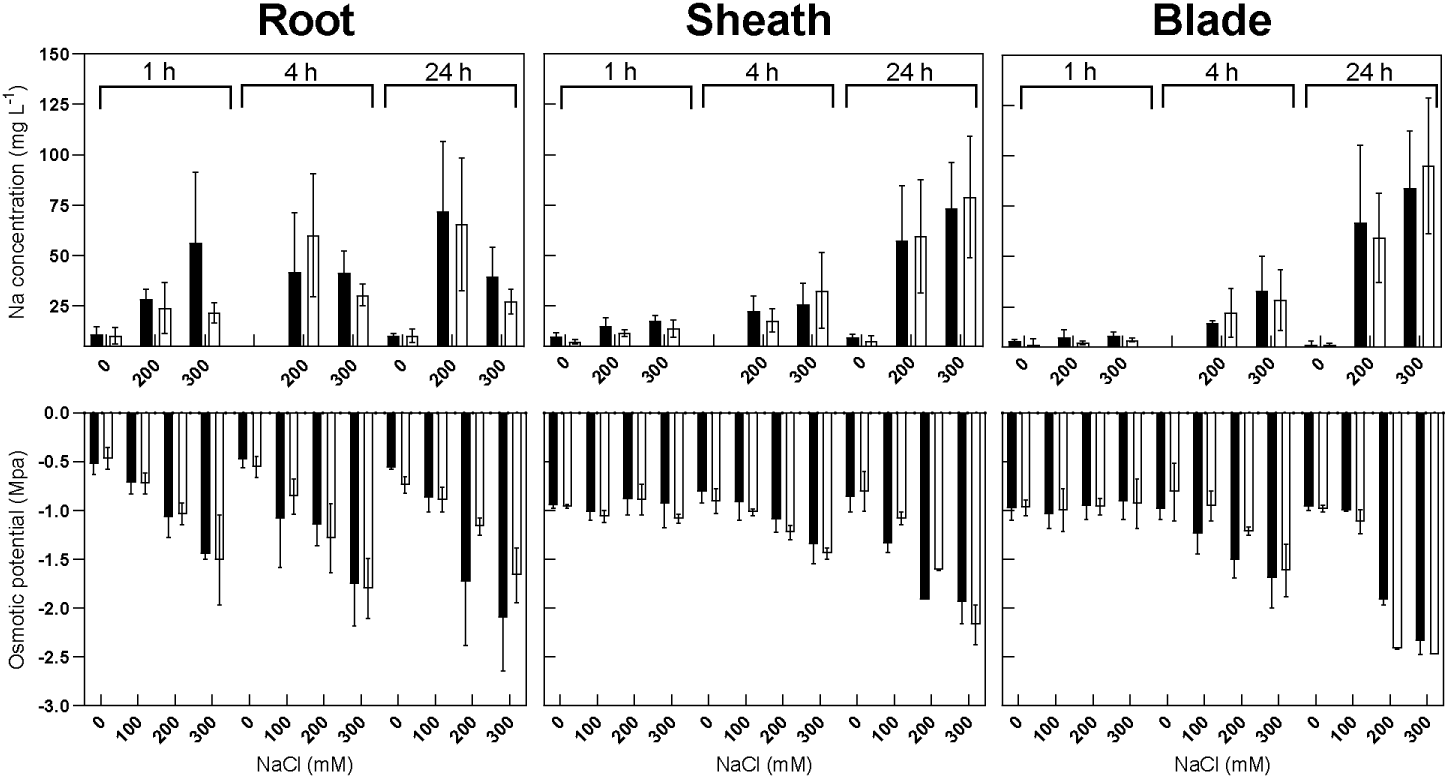
Sodium concentration (mg L^−1^) and osmotic potential (MPa) measured on root, sheath and blade of sorghum seedlings (10 days old) of two genotypes (RedLand B2 and IS9530). Seedlings were subjected to increasing sodium concentrations, 200 and 300 mM NaCl, during 1, 4 and 24 hours. Values are means ± SEM of three independent experiments for sodium concentration and three plants per genotype from four independent experiments for osmotic potential.

**Table 1.** Type III Analysis of Variance table with Satterthwaite’s method to evaluate the effect of genotype, treatment and time on sodium concentration (mg L^−1^) at each part of seedlings (root, sheath and blade). *** indicate significant differences (P<0.0001).

|  | Sum Sq | Mean Sq | NumDF | DenDF | F value | Pr(>F) |
| --- | --- | --- | --- | --- | --- | --- |
| **Genotype** | 221.2 | 221.2 | 1 | 114.04 | 0.8881 | 0.3480 |
| **Treatment** | 22306.9 | 11153.4 | 2 | 114.03 | 447.743 | **4.45E-12** *** |
| **Time** | 30872.9 | 15436.4 | 2 | 114.03 | 619.680 | **< 2.2e-16** *** |
| **Genotype:Treatment** | 5.4 | 2.7 | 2 | 114.02 | 0.0109 | 0.9892 |
| **Genotype:Time** | 128.1 | 64.0 | 2 | 114.06 | 0.2570 | 0.7738 |
| **Treatment:Time** | 13397.2 | 4465.7 | 3 | 114.04 | 179.272 | **1.34E-06** *** |
| **Genotype:Treatment:Time** | 218.0 | 72.7 | 3 | 114.01 | 0.2917 | 0.8313 |

Genotypes: RedLand B2; 9530 Treatment: 0; 200; 300 mM NaCl Time: 1; 4, 24 hours of treatment

Osmotic potential is composed of all the ions/solutes present along the entire plant (root, sheath and blade). Especially potassium is important in relation to sodium for plant metabolism and as indicator of salt tolerance (Munns and Tester, 2008). Low Na^+^/K^+^ ratio in the cytosol indicates that the species is salt tolerant as was reported for many species like Arabidopsis, barley, rice and wheat (Lu and Fricke, 2023). However, our results showed an opposite pattern (Table 2) indicating that another mechanism could be involved. To elucidate this, we investigated the structure and composition of microhairs and pricklets.

**Table 2:** Sodium potassium ratio of calculated for root, sheath and blade of sorghum seedlings (10 days old) of two genotypes (RedLand B2 and IS9530). Seedlings were subjected to increasing sodium concentrations, 200 and 300 mM NaCl, during 1, 4 and 24 hours. Values are means ± SEM of three independent experiments.

| Time (h) | Treatment (mM NaCl) | Part of seedling | Genotype |  |
| --- | --- | --- | --- | --- |
|  |  |  | RedLand B2 | IS9530 |
| 1 | 0 | Root | 0.40 ± 0.55 | 0.44 ± 0.26 |
| 1 | 200 | Root | 1.82 ± 0.29 | 2.69 ± 1.17 |
| 1 | 300 | Root | 10.07 ± 0.55 | 6.13 ± 1.06 |
| 4 | 200 | Root | 6.10 ± 0.96 | 1.65 ± 0.43 |
| 4 | 300 | Root | 1.48 ± 0.13 | 2.35 ± 1.09 |
| 24 | 0 | Root | 0.48 ± 0.09 | 0.47 ± 0.32 |
| 24 | 200 | Root | 4.23 ± 0.70 | 4.04 ± 1.37 |
| 24 | 300 | Root | 5.48 ± 2.47 | 2.06 ± 1.33 |
| 1 | 0 | Sheath | 0.34 ± 0.10 | 0.26 ± 0.19 |
| 1 | 200 | Sheath | 0.41 ± 0.63 | 0.46 ± 0.23 |
| 1 | 300 | Sheath | 0.56 ± 0.24 | 0.43 ± 0.5 |
| 4 | 200 | Sheath | 0.77 ± 0.20 | 0.68 ± 0.25 |
| 4 | 300 | Sheath | 0.81 ± 0.67 | 0.81 ± 0.81 |
| 24 | 0 | Sheath | 0.34 ± 0.35 | 0.51 ± 0.63 |
| 24 | 200 | Sheath | 2.56 ± 0.72 | 4.64 ± 0.81 |
| 24 | 300 | Sheath | 3.45 ± 0.69 | 6.04 ± 0.53 |
| 1 | 0 | Blade | 0.38 ± 0.15 | 0.20 ± 0.28 |
| 1 | 200 | Blade | 0.24 ± 0.54 | 0.26 ± 0.26 |
| 1 | 300 | Blade | 0.61 ± 0.34 | 0.40 ± 0.34 |
| 4 | 200 | Blade | 0.52 ± 0.11 | 0.78 ± 0.42 |
| 4 | 300 | Blade | 1.09 ± 0.68 | 0.70 ± 0.47 |
| 24 | 0 | Blade | 0.28 ± 0.53 | 0.28 ± 0.57 |
| 24 | 200 | Blade | 1.99 ± 0.54 | 2.96 ± 1.15 |
| 24 | 300 | Blade | 2.79 ± 0.51 | 2.65 ± 0.42 |

We identified two types of glandular hairs in sorghum leaves of both genotypes, pricklets and microhairs (Ceccoli *et al.,* 2015; McWhorter *et al.,* 1995). According to our hypothesis, we analyzed the ions content (mainly sodium) in these structures. The main result was that the sodium was absent in all analyzed hairs and leaves surface. Surprisingly, we detected the presence of silicon inside the pricklet at 300 mM NaCl after 24 h of treatment, but not in the microhair (Fig. 7).

**Fig. 7.**
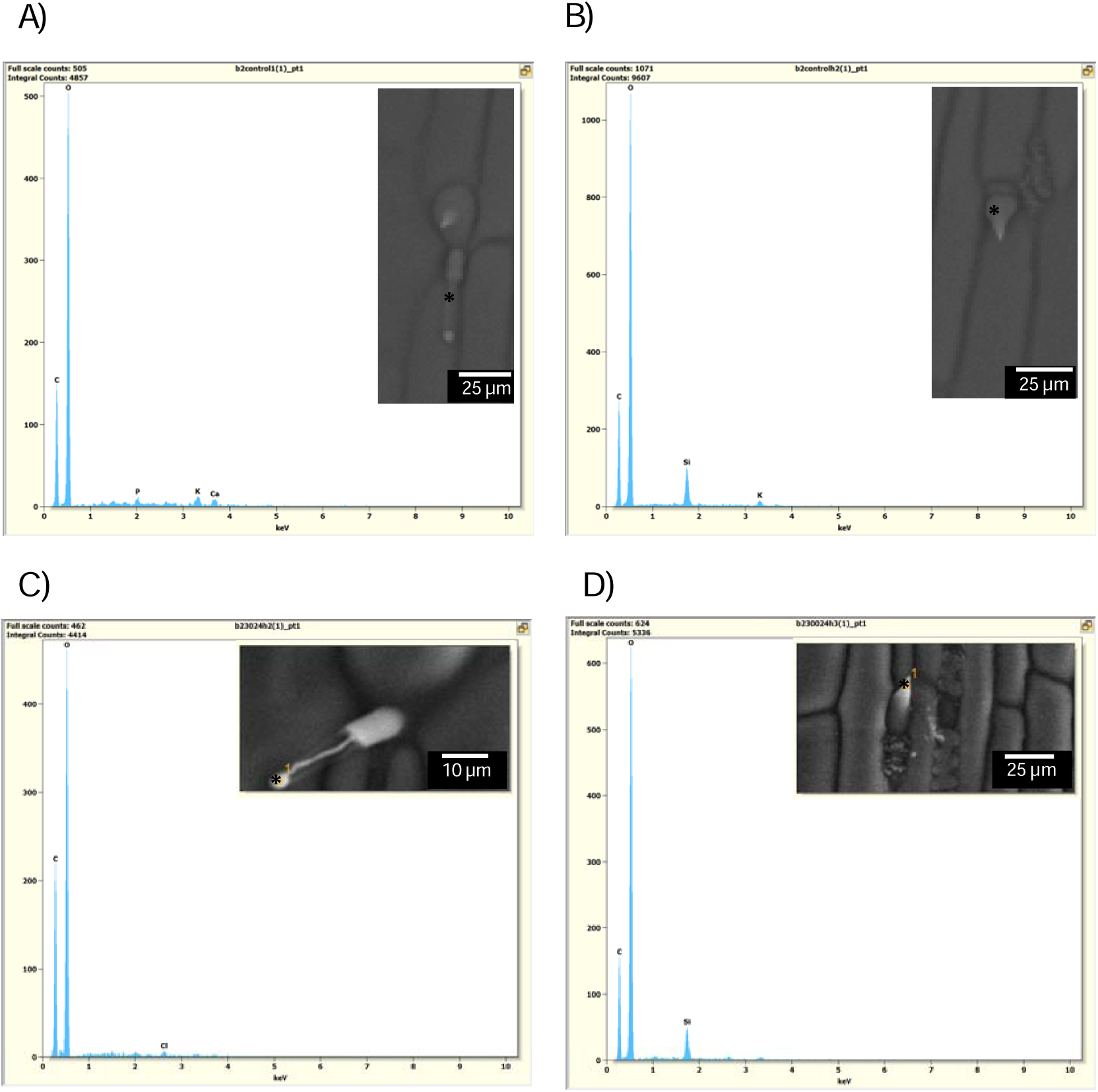
Photomicrographs and ion profiles of *S. bicolor* microhairs and pricklets from adaxial side of blades. Images were obtained from scanning electron microscopy (ESSEM, FEI Quanta 250). A) microhair and B) pricklet from control plants; C) microhair and D) pricklets treated with 300 mM NaCl for 24 hours. Asterisks in the picture indicate analysis place where ions were measured.

Thus, sorghum pricklets could be involved in salt tolerance at early development state. However, our results showed that pricklets internal composition did not include sodium, instead there was silicon (Fig. 7). Since salt is present in the soil solution at both 200 mM and 300 mM NaCl, the hypothesis is that Na^+^ would replace Si, for example in the cell walls. At 200 mM NaCl, it is seen in the root, sheath, and blade, while at 300 mM, it is mainly present in the aerial parts. Si would escape through the pricklets. At 300 mM, the excess sodium would break the Na-Si stoichiometry, and this would determine the release of Si to the leaf surface, while Na+ would be retained in the cell walls.

Several NIPs aquaporins transport silicon in various species such as rice, barley, maize and soybean (Handa *et al.,* 2022). It is well known that silicon is transported by LSI (Low Silicon 1) from soil to plant and the mechanisms are established (Mandlik *et al.,* 2020). In sorghum, the expression of two NIPs is known to be increased in salinity (Reddy *et al.,* 2015). These NIPs have a high degree of homology (98%) with those already reported suggesting that they could also transport silicon and participate in the response to salt (Fig. 8).

**Fig. 8.**
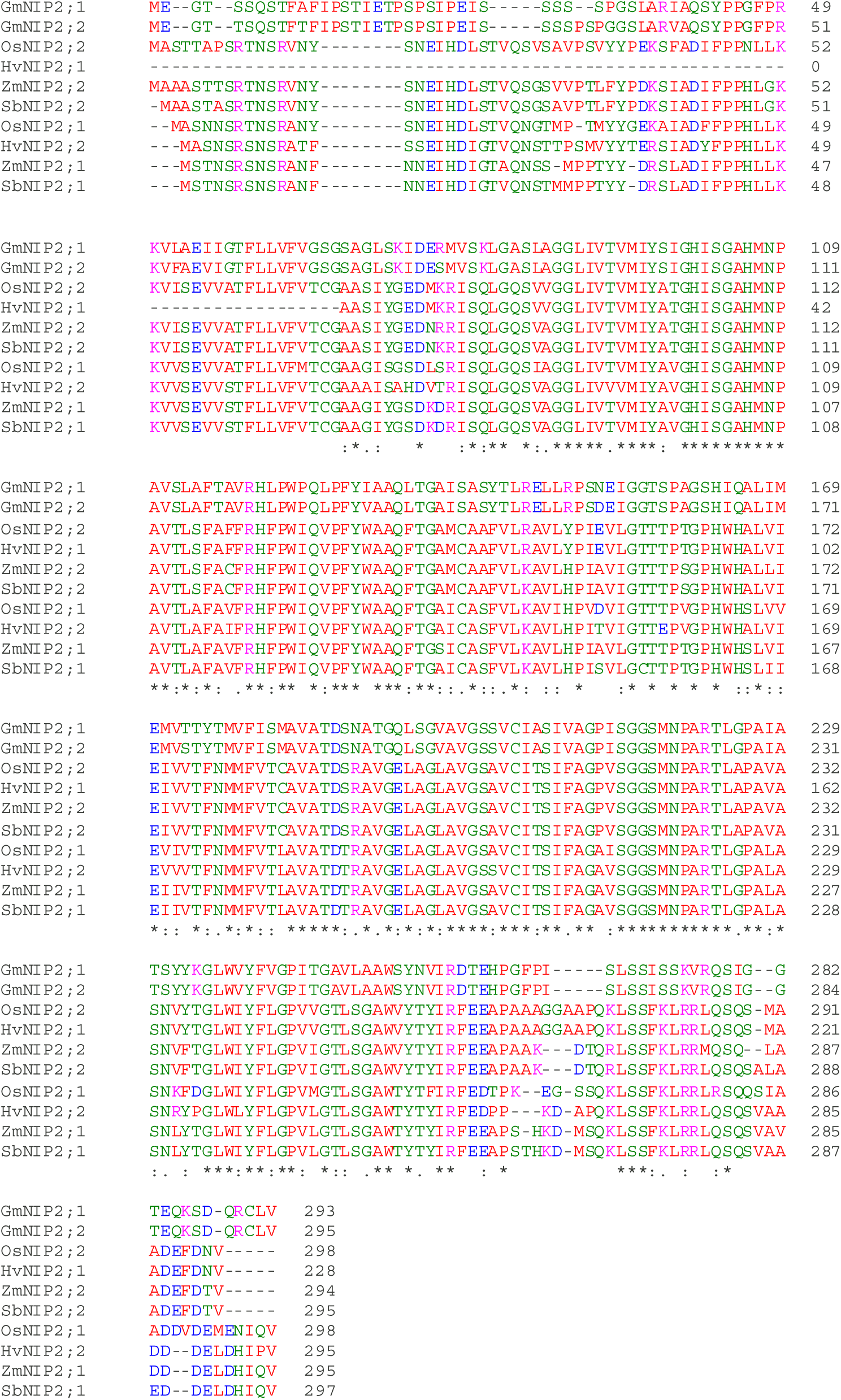
Sequence multiple alignments of NIPs aquaporin reported as silicon transporters by Handa *et al.,* (2022) and NIPs from *Sorghum bicolor*. Strictly conserved amino acids are marked with an asterisk and the amino acids of the same type are marked with two points.

Silicon is known to improve the response of plants to high salt concentrations (Peña-Calzada *et al.,* 2023; Mahmood *et al.,* 2025), although the mechanisms involved in this response are not yet fully understood. Some evidence shows that Si stimulates the synthesis of proline and cytokininin in cucumber (Zhu et. al., 2020), so the effect of “helping” the response to salt would be more due to an osmolyte issue. In short term salt stressed S*orghum bicolor* plants, Si application regulates PIP aquaporins activity and consequently root hydraulic conductivity was less affected (Liu *et al.,* 2015) However, we found that both sorghum genotypes would have a common mechanism that does not involve sodium extrusion but rather would use silicon to replace it in the cell walls and remove the sodium. Pricklets would be removing silicon instead of sodium. This new hypothesis is supported by our own results, where we demonstrated that the pricklets contain silicon instead of sodium.

Our work demonstrates that NaCl concentrations generate physiological changes in sorghum seedlings when they reach values over 100 mM. In this sense, at 200 mM NaCl the strategy would appear to be hydraulic redistribution throughout the plant, as demonstrated by the recovery of the RWC at 24 hours, given that the stomatal conductance and the hydraulic conductivity of roots remained low. However, at 300 mM NaCl the scenario is quite different. We have evidence to hypothesize that sorghum seedlings mobilize and accumulate Na^+^ in the cell walls, displacing silicon secreted by pricklets, since we observed the restoration of water balance after 24 hours. This hypothesis requires future experimental approaches to be verified.

## Supplementary data

Fig. S1. Time-lapse of sorghum seedling showing the phenotype throughout the salt concentration.

Fig. S2. Freehand cross-section photograph of sheath of *Sorghum bicolo*r seedlings.

## Acknowledgements

We would like to thank Marina Zanitti (from IBBEA CONICET) for providing access and assistance to flame spectrophotometer and Camila Dávila for edition of figure 2 and critical reading of the manuscript.

## Author contributions

MRS conceptualization, experimental design and execution; data curation and analysis, funding acquisition; PDC experimental execution; data analyses; MR and ASD experimental execution and technical support; MEM conceptualization, experimental design, execution and analysis; MRS and MEM wrote the manuscript.

